# Blood originates in hypoblasts during embryonic development

**DOI:** 10.1101/2025.11.07.687183

**Authors:** Yiming Chao, Hongji Li, Lu Liu, Zhengyi Tay, Shihui Zhang, Xiaolin Xu, Shao Xu, Yang Xiang, Degong Ruan, Yuanhua Huang, Guocheng Lan, Pengtao Liu, Ryohichi Sugimura

**Author notes:** Contributed equally (Yiming Chao, Hongji Li, Lu Liu).

## Abstract

Blood is essential for oxygen supply throughout life. The emergence of blood in the human embryo remains poorly understood. Our study leverages multiple stem cell embryo models and advanced lineage barcoding to unveil that hypoblast, originally regarded as forming the yolk sac wall, is heterogeneous and contributes to CDX2+ extraembryonic mesoderm, followed by hemoglobin+ cells as the first blood cells. CDX2 marks the hypoblast-to-hemoglobin+ cell trajectory that functionally sustains oxygen levels in embryo models. These hemoglobin+ cells molecularly and functionally resemble phagocytes. We show that the erythro-core regulatory network is poised in hypoblasts, and its boost endows erythropoiesis to both hypoblasts and phagocytes. Hypoblasts are the origin of the first blood in humans and non-human primates, providing a conceptual framework that earlier blood generation than expected fills the gap in the establishment of circulation. Further, the hypoblast is a place where primates may repurpose the phagocyte program to carry oxygen throughout embryos.

**Highlights:**

- Barcode-traced hypoblast fate in human embryo models;
- The first blood comes from the hypoblast that contributes to hemoglobin+ phagocyte-like cells;
- CDX2 marks hypoblast blood that sustains oxygen supply in embryos before heart formation;
- Erythro-core regulatory network endows erythropoiesis to human hypoblasts and phagocytes.

## Introduction

In mammals, blood cells are generated in successive waves, beginning with the yolk sac, from which nucleated erythrocytes and macrophages are generated, followed by the derivation of blood cells from multipotent hematopoietic stem and progenitor cells in the hemogenic endothelium of embryonic aorta-gonad-mesonephros ^1^. In mice, the extraembryonic mesoderm that contains progenitors of the earliest wave of blood cells is derived from the epiblast at early gastrulation ^2,3^. In contrast, in non-human primate embryos, extraembryonic mesoderm investing the primary yolk sac is present prior to gastrulation^4,5^. A lineage tracing study based on clonality of post-zygotic variants theorized that the hypoblast of the peri-implantation human embryo contributes to the yolk sac endoderm and extraembryonic mesoderm, with the latter participating in yolk sac hematopoiesis to generate the primitive blood cells ^6^, including blood cells that express embryonic hemoglobins ^7^. This hypothesis highlights a putative lineage trajectory from hypoblast, then extraembryonic mesoderm, to hemoglobin-expressing cells prior to gastrulation in the human and non-human primate embryo, and raises an intriguing evolutionary difference in blood cell ontogeny that primate hypoblast may serve as a complementary source of the first blood cell population.

The elucidation of the origin of blood cells in early human embryos has been achieved by computational analysis of single-cell and spatial datasets ^8,9^. Recent advances in stem cell-based embryoid bodies and organoid models offer an avenue for experimental study of the early developmental events ^10^ such as yolk sac hematopoiesis ^11–13^. In particular, the stem cell-based embryo model is acquiring a level of complexity close to the natural embryos ^14, 15^, such that yolk sac blood formation can be modeled in genetically modified models ^12^. Recent advances in transcriptomic and epigenomic profiles in single cells and molecular barcoding technology^16 17^ has further enabled tracking the hematopoiesis in human stem cell-based models to address the origin of the earliest population of blood cells.

We combined a cross-species primate embryo atlas, multiple stem cell embryo models, advanced lineage tracing, and CRISPRa exploitation of the core regulatory network. We showed that hypoblast contributes to erythropoiesis at peri-gastrulation. We identified that human and non-human primates develop CDX2+ extraembryonic mesoderm, then hemoglobin+ cells, before gastrulation. CDX2, originally identified as a tropho-ectoderm lineage determinator ^18^, but a study suggests its role in somitic mesoderm development after gastrulation ^19^. We showed that CDX2 was required for extraembryonic mesoderm commitment, and its loss led to hypoxic embryos. These hemoglobin+ cells mimic phagocytes with a rich EMT-signature, suggesting their locomotion in heatless embryos to supply oxygen. Our study demonstrated that the hypoblast is an alternative source to sustain embryo oxygen supply before the establishment of the heart and blood from gastrulation in human and non-human primates. We propose that the erythro-core regulatory network defined in this study repurposes phagocytes, the first immune cells invented ^20^, as an oxygen carrier.

## Results

### Human hypoblast comprises heterogeneous cell types and transitions to extraembryonic mesenchyme

Previously, we have shown that functional hematopoietic stem and progenitor cells can be generated from human pluripotent stem cells (hPSCs) through morphogen-directed differentiation of the hemogenic endothelium ^21^. In the present study, we exploited the extended lineage propensity for embryonic and extraembryonic cell types of the human expanded potential stem cells (hEPSCs) ^22^ (Supp. Fig. 1A) to generate hematopoiesis-competent embryoids (HCEB) that contain extraembryonic and embryonic hemogenic progenitors for elucidating the hematopoietic lineage trajectory in early human embryos. By Day 4 of in vitro development, the HCEBs showed the segregation of OCT4+, FOXA2+ and GATA6+ cells, suggesting the emergence of the epiblast and hypoblast (Supp. Fig. 1B). The presence of separate clusters of GATA6+ cells and FOXA2+ cells suggests that the hypoblast is a heterogeneous cell population. This was followed by the emergence of SOX2+ ectoderm, SOX17+ endoderm, and T+ mesoderm by Day 6, and that of TFAP2C+ NANOG+ SOX17+ primordial germ cells (PGC) by Day 8 (Fig. 1C; Supp. Fig. 1C, D). In Day 8 HCEBs the hypoblast, extraembryonic mesenchyme and CDX2: HBE-expressing cells were present (Fig. 1A). Of note, comparable populations of CDX2+ cells were present in the hypoblast and extraembryonic mesenchyme of human and Macaque embryos (Supp. Fig. 2). CD34+ hematopoietic progenitor cells and CD43+ and CD45+ blood cells were detected by Day 10 of culture (Fig. 1B) and here on to Day 30, an expanding population of blood cells was found among other cell types of the three germ layers and PGCs (Fig. 1C). In Day 10 HCEBs, amnion, hemogenic endothelium, and yolk sac (YS) endoderm cells were present (Fig. 1 D; Supp. Fig. 1C). Like blood-forming cardiac organoids ^23^, cardiac mesoderm cell were present in the HCEBs (Fig. 1C). Time-course scRNA-seq of HCEBs identified 3 cell type transitions in the hypoblast population similar to those previously shown in murine ^24^. The transitions were first to OCT4+ SOX17+ GATA6+ populations (17 --> 12), second to OCT4-lo SOX17+ GATA6+ FOXA2+ CDX2+ populations (10 --> 9 --> 8), third to SOX17 lo GATA6+ FOXA2+ CDX2+ hemoglobin+ (20 --> 11, 13, 16, 19) (Fig. 1E). In particular, the biphasic peak expression of SOX17 has been noted previously in PSC-derived yolk sac-like cell model ^25^ (Fig. 1F). The cell type transition progressed during the 8 days of in vitro differentiation (Supp Fig. 1E), with the enrichment of HBE1, HBG1, HBG2 and GYPA expression in Day 10 HCEBs (Fig. 1G). Taken together, human hypoblast is heterogeneous in cellular composition and progressively shifts toward CDX2+ cells.

**Figure 1.**
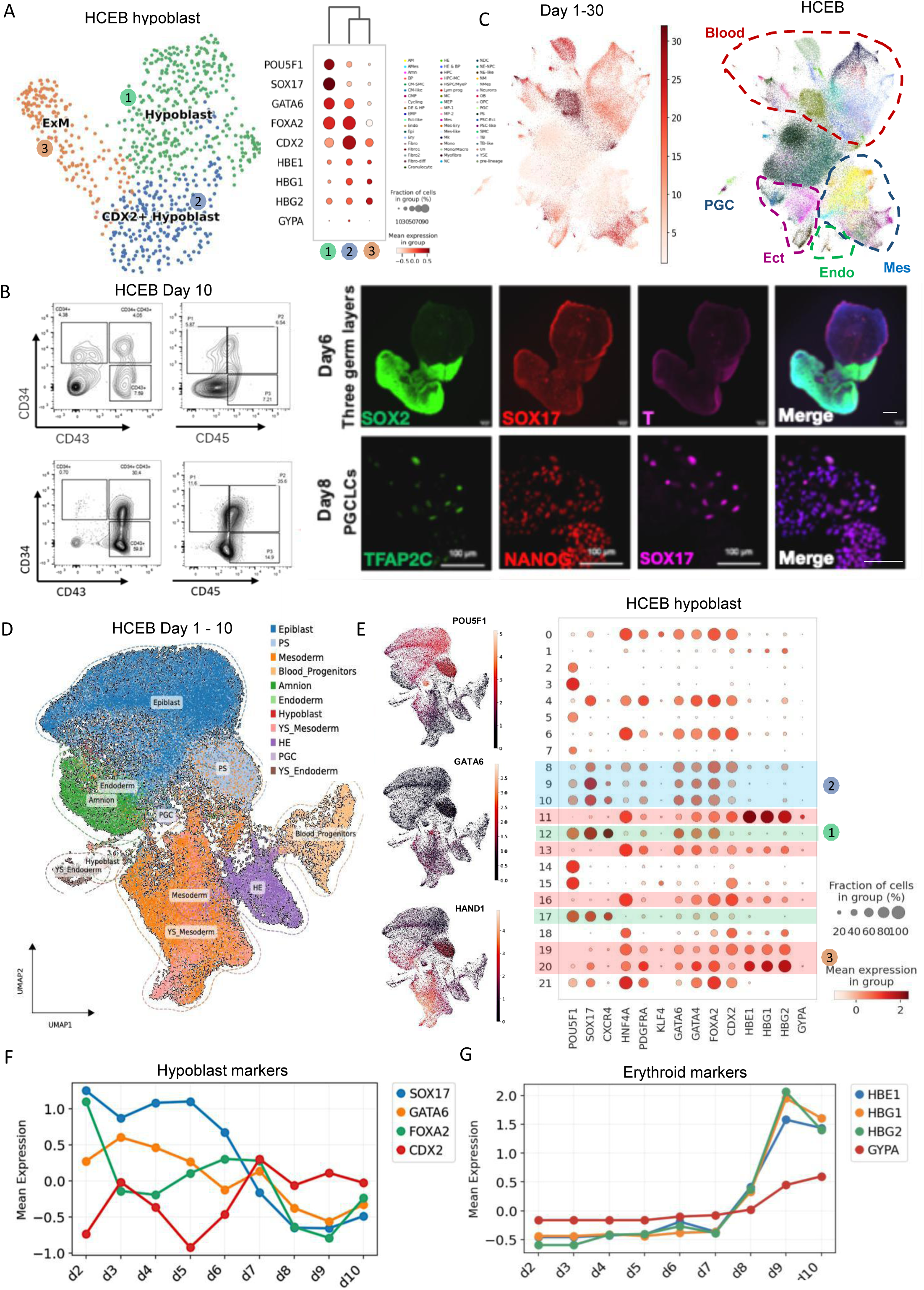
Human hypoblast comprises heterogeneous cell types and transitions to extraembryonic mesenchyme. A. scRNAseq of primary hypoblast, CDX2+ hypoblast, and extraembryonic mesenchyme (ExM) in Hematopoiesis-competent embryoid (HCEB) (left). Hypoblast markers SOX17, GATA6, FOXA2 expression were gradually expressed as well as CDX2 through trajectory from primary hypoblast, CDX2+ hypoblast to ExM (right). B. Hematopoietic markers CD34, CD43 and CD45 expression in Day 10 HCEB. C. Top: Hematopoietic lineage was expanded in HCEB up to Day 30. UMAP shows three germ layers, PGC, and blood cells were differentiated. Complete annotation is listed on the left. Bottom: Immunofluorescence of HCEB Day 6 and Day 8. Ectoderm (SOX2+), endoderm (SOX17+), mesoderm (T+) in Day 6 middle panel; PGC (TFAP2C+, NANOG+, SOX17+) in Day 8 lower panel. Scale bar = 100 um. D. UMAP of HCEB up to day 10 indicating blood, mesoderm, hypoblast, hemogenic endothelium, and yolk sac mesoderm. POU5F1, GATA6, and HAND1 expression were listed as indicators of epiblast and mesoderm. E. Hypoblast marker dot plot from HCEB. Hypoblast was heterogeneous. First was OCT4+ SOX17+ GATA6 populations (green: 17 --> 12), second was transition to OCT4 lo SOX17+ GATA6+ FOXA2+ CDX2+ populations (blue: 10 --> 9 --> 8), finally transition to SOX17 lo GATA6+ FOXA2+ CDX2+ hemoglobin+ (red: 20 --> 11, 13, 16, 19). F. Hypoblast markers in HCEB from time-course scRNA-seq. GATA6 expression wave was followed by FOXA2, then CDX2. SOX17 showed bi-phasic peaks. G. Erythroid markers in HCEB from time-course scRNA-seq. Hemoglobin genes (HBE1, HBG1, HBG2) and erythroblast marker GYPA appeared when CDX2 expression (Fig. 1F) plateau. hEPSC (human expanded potential stem cell); EB (embryoid body); HCEB (Hematopoiesis-competent embryoid); PS (primitive streak); YS (Yolk sac); HE (hemogenic endothelium); PGC (primordial germ cell); ExM (extraembryonic mesenchyme); CM (cardiac mesoderm); NE (neuroectoderm); DE (definitive endoderm); HP (hypoblast); BP (blood progenitors); EMP (erythro myeloid progenitor); Epi (epiblast); Ery (erythroblast); Endo (endoderm); Fibro (fibroblast); OB (osteoblast); OPC (oligo dendro progenitor); MP (myeloid progenitor); Mk (megakaryocyte); NC (neural crest); Mono (monocyte); Macro (macrophage); NM (nascent mesoderm); PSC (pluripotent stem cell); SMC (smooth muscle cell); TB (tropho-ectoderm); YSE (yolk sac endoderm).

We next examined the spectrum of hypoblast cell types in the peri-gastruloid model derived from the EPSCs (Fig. 2A). Segregation of OCT4+ epiblast and GATA6+ extraembryonic endoderm occurred at Day 2-4, with emergence of ectoderm, mesoderm and endoderm cell types at Day 8 (Fig. 2B). CD235A+ erythroids, comparable to those found in CS7-8 stage embryo and another peri-gastruloid model ^26^ also emerged at comparable stages (Fig. 2C). Like the HCEBs, distinctive subsets of cells expression GATA6, and FOX2A, were found in the hypoblast cluster (Fig. 2D). Time-series transcriptomics of embryonic endoderm populations revealed successive emergence of GATA6+/SOX17+ cells and FOXA2+ cells (Fig. 2E). Altogether, these data support that hypoblast comprises heterogenous cell types and transitions to extraembryonic mesenchyme.

**Figure 2.**
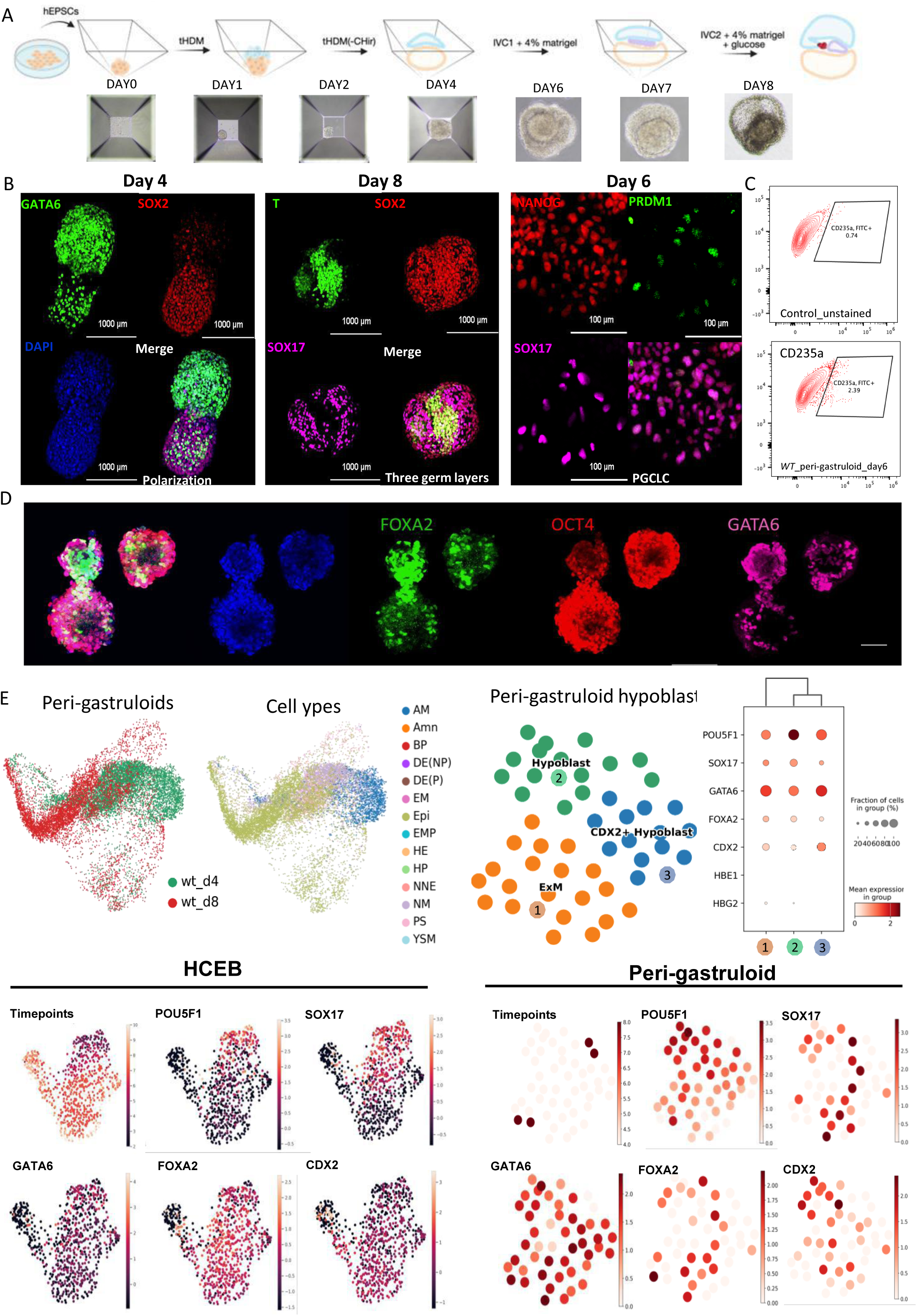
Peri-gastruloid hypoblast shows heterogeneity. A. Peri-gastruloid generation from human expanded potential stem cells (hEPSC). B. Epiblast (SOX2+)-hypoblast (GATA6+) polarization in the left panels Day 4; Ectoderm (SOX2+), endoderm (SOX17+), mesoderm (T+) in the middle panels Day 8; Primordial germ cells (NANOG+ PRDM1+ SOX17+) in the right panels Day 6. Scale bar = 1,000 um. C. Erythroblast marker CD235A expression in Day 8 peri-gastruloids. D. Hypoblast heterogeneity indicated by reciprocal expression of FOXA2 (inside layer of left peri-gastruloid) and GATA6 (outside layer of left peri-gastruloid). Scale bar = 1,000 um. E. Top left: A UMAP of peri-gastruloids at Day 8 was shown. Top right: Primary hypoblast, CDX2+ hypoblast, and ExM (extraembryonic mesenchyme) were shown on UMAP from Day 8 peri-gastruloids. Gradient expression of hypoblast markers, and CDX2 was detected in primary hypoblast, CDX2+ hypoblast and ExM. Bottom: Hypoblast of HCEB (left) and peri-gastruloid (right). SOX17+ GATA6+ cells were followed by FOXA2+ CDX2+ cells. ExM (extraembryonic mesenchyme). tHDM (N2B27 basal medium supplemented with 10 ng/mL FGF2, 10 ng/mL Activin-A, 1 μM CHIR99021 and 0.3 μM PD0325901); IVC1 ((76 mL Advanced DMEM/F12, 1 mL Glutamax (100x), 1 mL Insulin-Transferrin-Selenium-Ethanolamine (ITS-X) (100x), 10.8 μL β-estradiol stock (73.4 μM, final concentration 8 nM); IVC2 (identical to IVC1 except 20% FBS was replaced with 30% KSR). AM (advanced mesoderm); Amn (amnion); BP (blood progenitor); DE (NP) (definitive endoderm, not proliferating); DE (P) (definitive endoderm; proliferating); EM (emergent mesoderm); Epi (epiblast); EMP (erythro myeloid progenitor); HE (hemogenic endothelium); HP (hypoblast); NNE (nascent neuroectoderm); NM (nascent mesoderm); PS (primitive streak); YSM (yolk sa mesoderm).

### Hypoblast contributes to erythroblasts

We re-constructed the lineage trajectory of CDX2-positive cells using datasets of human and macaque embryos ^27^. CDX2 was found to be expressed in the hypoblast, yolk sac endoderm and extraembryonic mesoderm in both human and macaque fetal samples (Supp. Fig. 2). These findings suggest that in the EPSC-based HCEBs and peri-gastruloids, there may exist a branching of the hypoblast lineage to the extraembryonic mesenchyme separately (Supp. Fig. 3A). Notably, the expression of CDX2 in the third cell type is concordant with the upregulation of erythroid genes such as hemoglobin (HBE1, HBG1, HBG2) and CD235A/Glycophorin A (GYPA) (Fig.1F, G). At Day 4 of HCEB culture, which corresponds to the CS7 stage, two distinct erythroblast populations were present in the HCEBs. There were a major population of erythroblasts with enriched EMT signatures including genes of SNAI family and ZEB family were found, which was termed mesenchymal erythroblast (Mes-Ery), and a minor population of conventional erythroblasts (Ery2) (Fig 3A, B; Supp. Fig. 3B). Both erythroblast populations displayed BMP signaling signatures, which are also found in the hypoblast, suggesting these erythroblasts may have a lineage relationship with the hypoblast.

**Figure 3.**
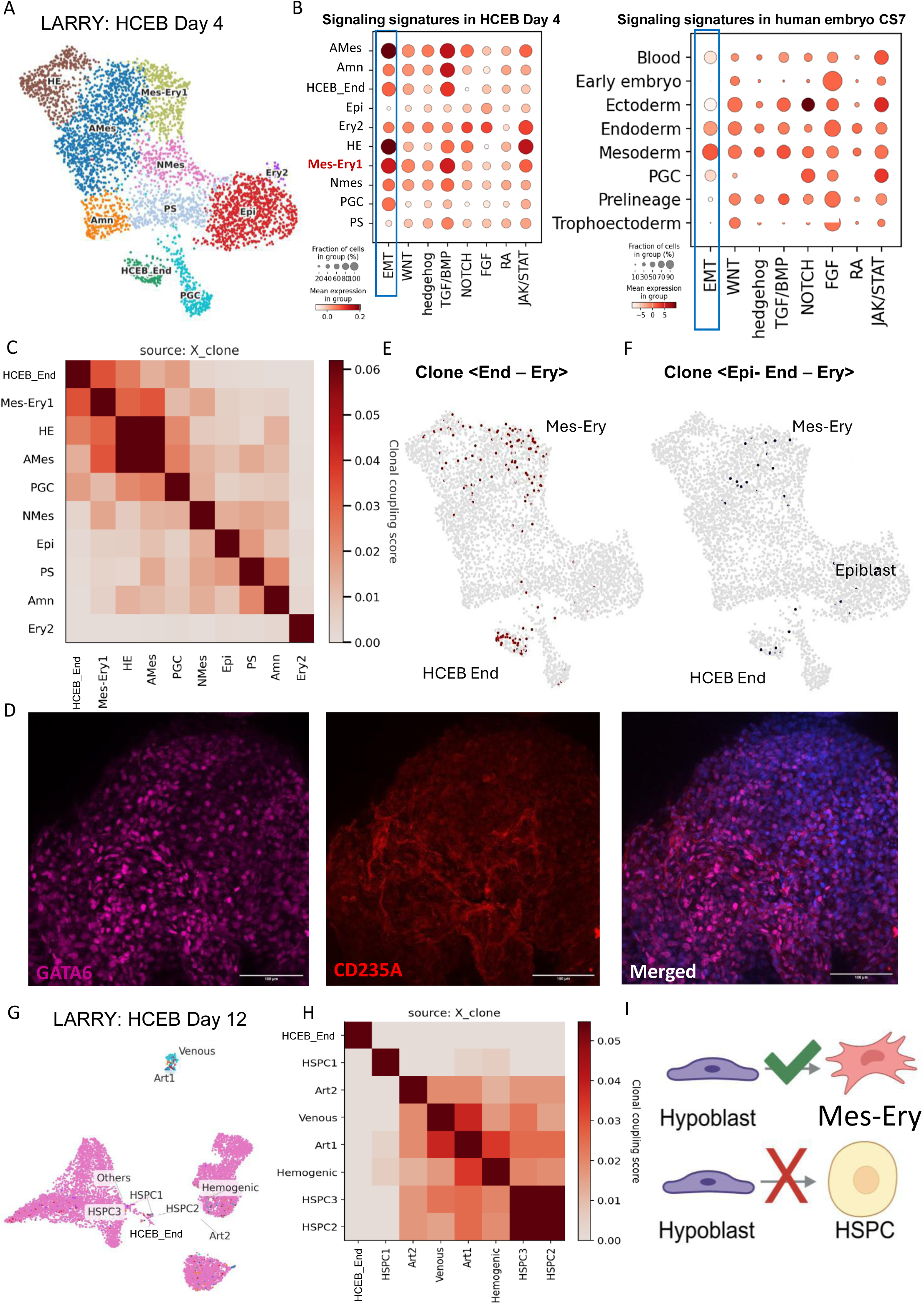
Hypoblast contributes to erythroblasts. A. UMAP of Day 4 HCEB showing erythroblast populations including Mes-Ery and Ery. HCEB endoderm comprised definitive endoderm and hypoblast derivative. B. EMT signature was rich in erythroblasts in both Day 4 HCEB (left) and CS7 human embryo (right). Erythroblasts comprised 2 populations. One was mesenchymal erythroblast (Mes-Ery) with a high EMT signature. The other one is conventional erythroblasts (Ery). C. X_Clone analysis of LARRY barcode sharing among each cell population. HCEB endoderm most commonly shared barcodes with Mes-Ery, indicating they came from the same lineage. D. Immunofluorescence showed that GATA6+ hypoblasts were co-localized with CD235A+ erythroblasts in Day 4 HCEB. Scale bar = 100 um. E. Hypoblast derivative to erythroblast contribution was major. LARRY clone detection showed that the majority of Mes-Ery was contributed by HCEB endoderm. F. Definitive endoderm contribution to erythroblasts was a few. LARRY clone detection showed that a tiny fraction of epiblast contributed to HCEB endoderm and Mes-Ery. G. UMAP of Day 12 HCEB showing hypoblast, endothelium, hemogenic endothelium and HSPC. H. X_Clone analysis of LARRY barcode from Day 12 HCEB with each cell population. Hypoblast did not share barcodes with hemogenic endothelium (HE) nor HSPC. I. LARRY analysis indicated that hypoblasts committed to Mes-Ery, but not HSPC. HCEB (Hematopoiesis-competent embryoid); End (Endoderm); Mes-Ery (mesenchymal erythroblast); Ery (erythroblast); HE (hemogenic endothelium); Ames (Advanced mesoderm); PGC (primordial germ cell); NMes (Nascent mesoderm); Epi (epiblast); PS (primitive streak); Amn (amnion); HSPC (hematopoietic stem and progenitor cell); Art (artery).

To validate the lineage trajectory inferred from transcriptome data, we traced the cell lineage experimentally by molecular barcoding cells of HCEBs (Supp. Fig. 3C) and analyzing the clonal relationship of the cellular descendants in the developing HCBEs. A total of 2,626 clones containing LARRY barcodes were recovered, with good coverage of cell types in the HCEB (Supp. Fig. 3D, E). Significant pairing was observed between HCEB endoderm, constituted of definitive endoderm and descendants of hypoblast, and the Mes-Ery cells (Fig. 3C). The pairing of hypoblast and Mes-Ery cells was commensurate with the co-localization of GATA6+ hypoblast cells and with CD235A+ erythroblasts in HCEB (Fig. 3D). We further interrogated whether definitive endoderm may contribute to Mes-Ery. Notably, while the majority of Mes-Ery was contributed by HCEB endoderm (Fig. 3E), the epiblast-derivative (e.g., the definitive endoderm) contributed to only a minor fraction of Mes-Ery cell (Fig. 3F). Further, the hemogenic endothelium or the hematopoietic stem and progenitor cells (HSPC) (Fig. 3G) did not share any clones with the hypoblast (Fig. 3H), suggesting that the hypoblast contributes to the Mes-Ery cells but not the HSPC (Fig. 3I). These findings suggest that the hypoblast is a significant source of mesenchymal erythroblasts whereas the epiblast-derived definitive endoderm in the HCEB endoderm ^28^ makes a minor contribution. Altogether, these data support the concept that the hypoblast contributes to erythroblasts.

### Functionality of mesenchymal erythroblast

In the mouse, CDX2 cooperates with EOMES in cell fate determination in trophectoderm ^29^, and the development of the allantois that contributes to extraembryonic mesoderm and yolk sac hematopoiesis ^30^. We hypothesized that CDX2 marks the human hypoblast to hemoglobin+ cell trajectory and might regulate this process of extraembryonic erythropoiesis. Consistent with enhanced CD235A expression, peri-gastruloids generated hemoglobin epsilon (HBE1)+ cells, suggesting the embryonic form of hemoglobin delivers the oxygen carrier attribute ^31^ (Figure 4A). CDX2 -/- peri-gastruloids displayed raised pimonidazole signal, indicating they were experiencing hypoxia (Figure 4B-C). CDX2 -/- peri-gastruloids were predominantly comprised of epiblast and less hypoblast, suggesting a role of CDX2 in hypoblast development (Figure 4D, E). GATA6+ cells persisted, but FOXA2+ cells were reduced, and SOX17+ cells were missing (Figure 4F). CDX2-/- peri-gastruloids downregulated EMT signature, suggesting that the population of EMT-active (mesenchymal) erythroblasts may be reduced. Metabolic pathways, such as glycolysis and OXPHOS, were downregulated, further supporting the occurrence of hypoxia (Figure 4G). The findings indicate that CDX2 activity underpins the cellular heterogeneity of the hypoblast, and loss of CDX2 function leads to hypoxia in peri-gastruloids.

**Figure 4.**
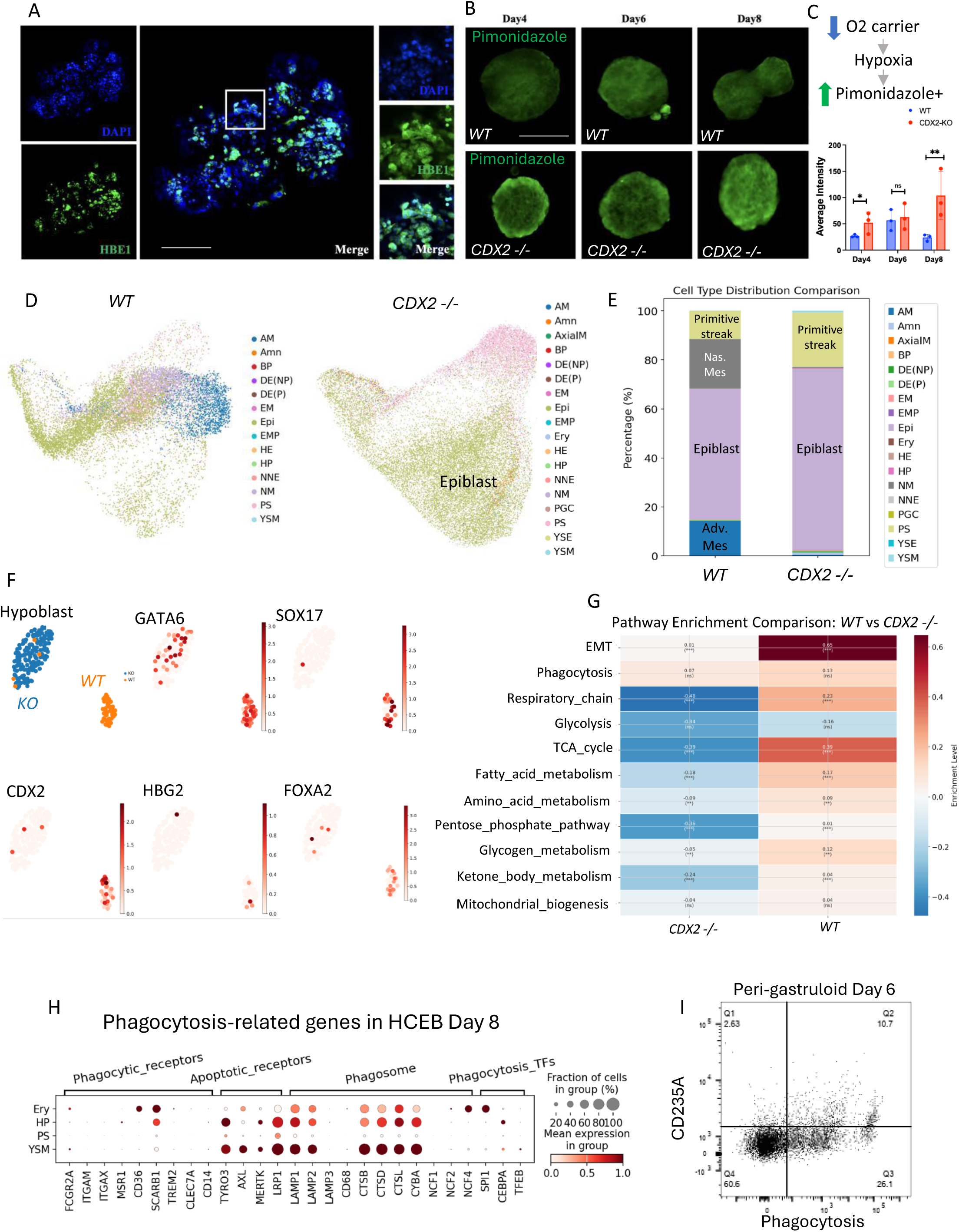
Functionality of mesenchymal erythroblast. A. Immunofluorescence of hEPSC-derived peri-gastruloid at Day 6. Peri-gastruloid expressed embryonic hemoglobin epsilon (HBE1). Scale bar = 200 um. B. Hypoxic indicator pimonidazole increased in *CDX2 -/-* peri-gastruloids. Immunofluorescence of pimonidazole in Wt (upper panels) and CDX2 -/- (lower panels) at Day 4, Day 6 and Day 8. Scale bar = 600 um. C. Schematics of how CDX2 deletion led to hypoxic peri-gastruloids. Quantification of fluorescence intensity of pimonidazole between Wt and CDX2 -/- peri-gastruloids. N=3 for each experiment. Data were shown with an error bar SEM. * p value <0.05. ** p value <0.01. D. UMAP of *WT* vs *CDX2 -/-* peri-gastruloid at Day 8. AM (Advansed mesoderm); Amn (amnion); BP (blood progenitor); DE (NP) (definitive endoderm not proliferating); DE (P) (definitive endoderm proliferating); EM (emerging mesoderm); Epi (epiblast); EMP (erythro myeloid progenitor); HE (hemogenic endothelium); HP (hypoblast); NNE (nascent neuroectoderm); NM (nascent mesoderm); PS (primitive streak); YSM (yolk sac mesoderm). E. Cellular composition of *WT* vs *CDX2 -/-* at Day 8 peri-gastruloid. Epiblast predominated *CDX2 -/-* peri-gastruloids. F. UMAP of hypoblast population in peri-gastruloids. *CDX2 -/-* peri-gastruloids lost SOX17+ hypoblasts. FOXA2 and HBG2 were reduced to diminished. G. GSEA of EMT, glycolysis and OXPHOS pathways were downregulated in WT vs *CDX2 -/-* peri-gastruloids. H. Dotplot of Day 8 HCEB. Hypoblasts, extraembryonic mesodermal cells, and erythroblasts expressed phagocyte signatures. I. Bead-based phagocytosis assay of Day 8 peri-gastruloids by flow cytometry readout. CD235A+ erythroblasts in peri-gastruloids possessed functional phagocytosis. Ery (erythroblast); HP (hypoblast); PS (primitive streak); YSM (yolk sac mesoderm).

Peri-gastruloids correspond to CS7-8 human embryos ^26^, in which circulation is not yet established. That this population of Mes-Ery cells displayed EMT phenotype and cellular attributes of phagocytes, suggests that they may behave like motile cells. The locomotive capacity of Mes-Ery cells may enable delivering oxygen carried by these cells to the tissues in an embryo that has yet to establish the circulatory system. We further examined whether hypoblast-derived erythroblast mimics the phagocytes, one of the most ancient blood cell types ^32^. The phagocyte program was expressed in hypoblast, extraembryonic mesodermal cells, and most abundantly in erythroblast (Figure 4H). Notably, CD235A+ erythroblast in peri-gastruloids demonstrated functional phagocytosis (Figure 4I). These findings led to the hypothesis that hypoblasts may exploit the phagocyte program to deliver oxygen in the early human embryo.

### Core regulatory network for hypoblast erythropoiesis

We next tested if activation of the erythro-core regulatory network may drive hypoblast erythropoiesis in the extraembryonic endoderm (hXEN) derived from human EPSCs ^33^ (Supp. Fig. 4A). EPSC-derived hXEN was identified by expression of GATA6, SOX17 and PDGFRA and lack of expression of OCT4, SOX2, and NANOG, and low level of SSEA4 expression (Supp Fig 4B-D), and the XEN genes are enriched in the hypoblast and extraembryonic mesenchyme (Supp Fig 4E). Culturing hXEN in hemogenic media led to the emergence by Day7 of CD34+ and CD43+ hematopoietic progenitors (Fig 5A). We examined the function of 30 genes of the core regulatory networks of erythropoiesis in human and macaque fetuses separately ^34^ (Supp Fig 4F, G) by CRISPRa-activation of these genes in hXEN cells. In view of the aforementioned finding that the Mes-Ery cells display molecular phenotype reminiscent of that of phagocytes, we include candidate genes that may promote erythroblast marker expression in human phagocytes in the pool library. We examined whether these genes promotes hypoblast erythropoiesis. Taking its resemblance to phagocytes as shown above, we triaged candidate genes that endow erythroid features to phagocytes. We conducted CRISPRa-activation ^35^ triage of 30 genes in a pooled library to promote erythroblast marker CD71 expression in human phagocytes (Figure 5B). The activation of 30 genes in a pool increased CD71 level in THP1 phagocytes (Figure 5C). We further validated that these genes converted hXEN to the erythroid lineage (Figure 5D). Altogether, we defined the erythro-core regulatory network that upregulated erythropoiesis from both hypoblasts and phagocytes. This regulatory network also endowed both phagocytes and hypoblasts with hemoglobin expression, suggesting that erythropoietic activity in the hypoblast may repurpose another blood cell type, the phagocytes, to motile erythrocyte-like cells in peri-implantation and immediate post-implantation primate embryos

**Figure 5.**
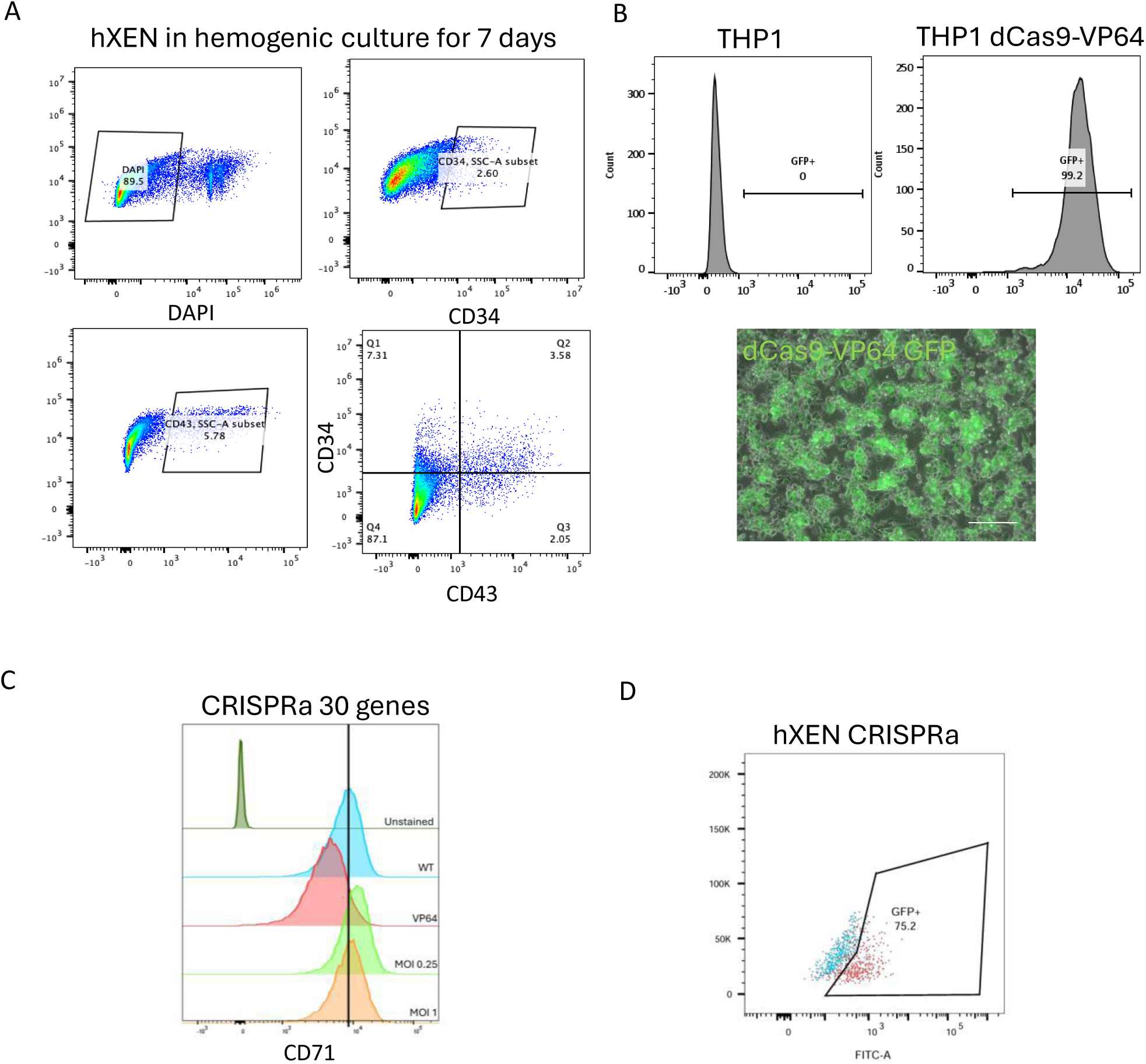
Core regulatory network for hypoblast erythropoiesis. A. Flow cytometry of hXEN expressed hematopoietic markers CD34 and CD43 upon exposure to hematopoietic cytokines and Activin A culture for 7 days. B. Generation of THP1 monocyte expressing dCas9-VP64 indicated by GFP. Fluorescence microscope showed GFP+ THP1 expressing dCas9-VP64. Scale bar = 100 um. C. CD71 erythroblast marker was up in CRISPRa-pooled gRNA-induced THP1 monocytes. Compared with VP64 that downregulated CD71, addition of pooled gRNA (MOI 0.25 and 1.0, respectively) rescued the CD71 level than WT. D. hXEN expressing dCAS9-VP64 and gRNAs. hXEN (human extraembryonic endoderm).

## Discussion

The fundamental question in blood development is, what is the first appearance of blood in life? Though the yolk sac is recognized as the primitive hematopoiesis, its origin was puzzling in human embryos. The existence of extraembryonic mesodermal cells around the primary yolk sac, way before gastrulation starts, questioned their origin. This study determined that hypoblasts contribute to CDX2+ extraembryonic mesoderm, subsequently to blood. The recent study suggested that the epiblast contributes to secondary hypoblast ^28^. Whether the first blood comes from primary or secondary hypoblasts will be the next question to be addressed. The physiological relevance of the first blood coming from the hypoblast is to maintain oxygen supply in the embryos before heart formation. The loss of CDX2 resulted in hypoxic peri-gastruloids. CDX2 is originally identified as tropho-ectoderm specifier ^18^. As tropho-ectoderm is lacking in peri-gastruloids ^26^ unlike other complete embryo models ^36–40^, CDX2 roles in our study would be primarily in extraembryonic mesodermal cells. How CDX2 regulates hypoblast erythropoiesis is a fundamental question. One possible explanation is that CDX2 might inhibit SOX2 and limit their commitment in epiblast, as shown in CDX2 -/- peri-gastruloid became predominantly epiblasts in our study, and CDX2 is a lineage specific transcriptional suppressor in another report ^41^.

The intriguing observation we made is that hypoblast erythropoiesis, both molecularly and functionally, resembled phagocytes. Considering phagocytes are locomotive cells in the tissues, this trait might be what oxygen-carrying cells need before the heart beat starts. Indeed, the same core regulatory network we identified was able to endow erythropoiesis to both phagocytes and hypoblasts. Phagocytes are the oldest blood type in evolution ^32^. Zebrafish use phagocytes to select true hematopoietic stem cells from hemogenic endothelium ^42^. We speculate that primates repurposed a phagocyte program for hypoblast to carry oxygen.

We leveraged two different human embryo models combined with molecular barcoding. HCEB is capable of hematopoiesis and advantageous to look at blood lineage. Peri-gastruloid is precise in morphology. We got consistent results in these two embryo models in hypoblast heterogeneity and their fate in blood. We further backed up our claims by cross-referencing the human and macaque fetus dataset. LARRY molecular barcoding is suitable for scRNAseq readout and allows enough diversity of barcodes, and is employed in dissecting hematopoietic stem cell fate ^43,44^.

The limitations of this study are as follows. Our barcode was based on the revised stem cell model that the early stages of stem cells are fate-committed. Although we observed that each stem cell has a fate bias, and some stem cells showed commitment to hypoblast and blood, to further identify the lineage trajectory, directly barcoding hXEN is needed. The hypothesis that hypoblast employs a phagocyte program to carry oxygen in heartless embryo is speculative, and live imaging of hemoglobin+ cells in embryo models will be required.

## Methods

### Stem cell line culture and maintenance

Human expanded potential stem cells (hEPSCs) developed by Pentao Liu’s lab ^45^ were used in this study. The lines C5 and EM1 are registered in the Centre for Translational Stem Cell Biology for research use. All cell lines were cultured at 37 °C in a 5% CO2 humidified incubator with daily media changes. Cells were periodically examined for Mycoplasma contamination via PCR. Cell lines were authenticated by genomic PCR, immunostaining, and in vitro differentiation tests.

hEPSCs were maintained in condition as described previously ^22^. Cells were maintained in hEPSC medium, on irradiated SNL plates that were pre-coated with 0.2% gelatin. Medium was changed daily until 70% confluency. hEPSCs were dissociated into single cells with TrypLE at 37°C for 3-5 minutes and passaged to a new SNL plate at a 1:10 - 1:20 ratio every 3-4 days for routine passaging.

We followed ISSCR ethical regulation of human embryo model research ^46^. IRB is submitted to HKU.

### HCEB generation

Hematopoiesis-competent embryoid (HCEB) was generated according to previous study^47^. In brief, hEPSCs were maintained in chemically defined medium in good condition for the following differentiation ^22^. The hEPSC medium was removed and changed to KSR medium for 3 days differentiation. Cells were then digested using TrypLE (12605036; Gibco) at 37°C for 5 minutes. After removing TrypLE, feeder-culturing medium was added to the cells, which were harvested carefully to avoid harsh pipetting.

Cells were centrifuged at 300g for 3 minutes, and the supernatant was gently aspirated. Cells were resuspended in KoSR medium supplemented with 10 μM ROCK inhibitor Y-27632. Hanging drops of 30 μl each, containing 5,000 cells per drop, were placed on the inner surface of the lid of a 100-mm petri dish (20100; SPL), with approximately 30 drops per lid. The dish was filled with PBS to maintain humidity, and the lid was inverted and placed over the dish. The dishes with EBs were incubated for 3 days.

3 days after, all EBs were collected with a 1,000-μl pipette and centrifuged at 20g for 1 min. The supernatant was carefully removed. EBs were cultured in an ultra-low-attachment plate with STEMdiff A medium (STEMdiff Hematopoietic Kit; STEMCELL Technologies) for 3 d followed by STEMdiff B medium (STEMdiff Hematopoietic Kit; STEMCELL Technologies) for 7 d as described previously. Successful generation of HCEB was benchmarked by elongation of embryoid as shown in other studies ^48,49^.

### Peri-gastruloid generation

Peri-gastruloids were generated following the protocol described previously ^26^. Briefly, hEPSCs were enzymatically dissociated into single cells using TrypLE at 37°C for 3-5 minutes, then resuspended in tHDM supplemented with CEPT. After digestion, cells were incubated sequentially on 0.1% gelatin-coated plates for 2 hours at 37°C to remove most of SNL feeder cells. After filtration through a 40 μm strainer, cells were counted and seeded into AggreWell800 plates pre-treated with anti-adherence solution at 45–90 cells per microwell in tHDM. Plates were centrifuged at 120g, 3min to facilitate cell aggregation. CEPT was added into tHDM on day 0. Half medium change was done on day 1 (tHDM without CEPT), followed by full medium changes on days 2 and 4 with different media respectively. No medium change was performed on day 5. On day 6, peri-gastruloids exhibiting characteristic embryonic disc-like morphology and yolk sac cavities were selected under microscopy and transferred to low-attachment plates for further culture. Due to the specific aims of our study, cultures were maintained only until day 8.

### CDX2 -/- human EPSC generation

CDX2 -/- hEPSC line was kindly provided by Pengtao Liu lab. Briefly, the ORF was deleted by Cas9/RNP electropolation followed by clone pick up, and genotyping by Sanger-seq to identify homozygous knock out lines.

### hXEN induction and hematopoietic differentiation

For differentiation of hEPSCs toward epiblast and hypoblast-like cells, cells were plated on SNL-coated plates using LCDM ^33^. After hEPSCs confluency reached 60%, the medium was switched to FTW, consisting of N2B27 in DMEM/F12 basal medium supplemented with 20 ng/mL FGF2, 20 ng/mL Activin-A, and 1 mM CHIR99021. Daily medium changes with fresh FTW were performed over the subsequent three days. The differentiated cells were evaluated on day 4 via immunostaining to analyze the co-cultured epiblast- and hypoblast-like cells. The mixture of differentiated cells was amenable to maintenance and passage.

### CRISPRa in hXEN

The gRNA sequences for 30 genes were selected from the CRISPRa Library (Addgene #60956), with 5 gRNAs for each gene. Pooled sgRNAs were ordered from Integrated DNA Technologies at 100 pmol scale (ordered as ATCTTGTGGAAAGGACGAAACACCG- [20bp protospacer]- GTTTTAGAGCTAGAAATAGCAAGTT).

70-bp oligomers were cloned into the CROPseq-Guide-Puro-EGFP vector (modified from Addgene plasmid #86708) using Gibson assembly as described ^50^. Cloned libraries were then sequence-verified by next-generation sequencing.

Lentiviral libraries were transduced into hXEN differentiated from the dCas9-VP64 EPSCs line at a multiplicity of infection (MOI) below 0.3. After 7 days of culture, cells were collected and subjected to bulk RNA sequencing.

### Flow cytometry

For direct cell surface stains, cells were spun at 300 x g for 3 minutes and washed twice with FACS buffer (PBS + 2% FBS). Cells were resuspended in 100 μL of FACS buffer containing 5ul FcX (human, Biolegend 422302) and incubated at room temperature for 5 minutes. Cells were then stained with conjugated antibodies in 100 uL of FACS buffer for 30 minutes on ice in the dark. Following incubation, cells were washed twice in FACS buffer and resuspended with DAPI buffer at an appropriate density for analysis or sorting. Data were acquired on a BD LSR Fortessa flow cytometer (BD Bioscience) and analyzed using the FlowJo (Flowjo 10.4, LLC) software.

### Immunofluorescence staining

Samples, including regular attached cell culture and peri-gastruloids at various stages, were washed with PBS before fixation with 4% paraformaldehyde (PFA) at room temperature for 15 min or at 4°C overnight. Fixed samples were then washed with PBS three times followed by permeabilization with 0.2% Triton X-100 in PBS at room temperature for 15 min. Subsequently, samples were washed 2 times with PBS and then blocked with 5% Bovine Albumin Fraction V (BSA) in PBS at room temperature for 1-4 h. Primary antibody dilution was prepared in 1% BSA in PBS. Samples were incubated in the primary antibody dilution at room temperature for 2 h or at 4 °C overnight followed by three PBS washes for 5min each. The secondary antibodies were diluted in 1% BSA in PBS and incubated with the samples for 1 h at room temperature away from light, followed by three washes in PBS for 5 min each. Samples were then stained with diluted DAPI in PBS for 15 minutes away from light, followed by three washes in PBS for 1 min each. Samples were then mounted using ProLong Glass Antifade (Invitrogen).

### Image acquisition and processing

Attached samples were cultured and imaged on 8-well μ-slides (ibidi). Peri-gastruloids were placed in 96-well glass-bottom black plates, µClear (Greiner) for imaging. Images were acquired using a Nikon Eclipse Ti2-E confocal microscope and processed using Fiji/ImageJ software.

### Lentiviral production

24 h before transfection, HEK293T cells were seeded in 100mm culture dishes at a density of 7-9e6 cells per well in 10 mL of DMEM + 10% FBS + 1% penicillin-streptomycin. The next day, cells were transfected using polyethylenimine (Polysciences) at a 5:1 PEI:plasmid ratio. Briefly, Opti-MEM and PEI were combined with a DNA mixture of the packaging plasmid pCMV_VSVG (Addgene plasmid no. 8454), psPax2 (Addgene plasmid no. 12260), and the transgene (CRISPRa library). The transfection mix was vortexed for 1min and incubated at room temperature for 15 min. After incubation, 1ml of transfection mix was added dropwise to the surface of the HEK293T cells. Dishes were then transferred to a 37 °C incubator for 8-12 h, after which the media was removed and replaced with DMEM +5% FBS + 1% penicillin-streptomycin.

Lentiviral-containing supernatants were collected 48 and 72 hours post-transfection and filtered through a 0.45 µm filter (Merck). Finally, the virus was concentrated 100-fold through ultracentrifugation. The 50mL tubes were centrifuged at 20,000 × g, 4°C for 2h, and the pellet was resuspended in Opti-MEM, aliquoted, and stored at -80 °C until further usage.

### Lentiviral titering

To determine lentiviral titer for transductions of pooled CRISPRa libraries and LARRY, THP1 cell line was transduced in 96-well plates with a range of virus volumes (e.g., 12, 2.4, 0.48, and 0.096 uL virus) with 1e5 cells per well. The plates were centrifuged at 1000 × g for 2 h at 33 °C. Three to four days post-transduction, cells were analyzed for fluorescence marker using flow cytometry on a BD LSRFortessa. A viral dose resulting in around 30% transduction efficiency, corresponding to an MOI of 0.3, was used for subsequent library screening and LARRY barcoding.

### Lentiviral barcoding using the LARRY system

Construction of lentiviral pLARRY vectors. pLARRY-eGFP vectors were obtained from Addgene plasmid 140025. The plasmids were packaged in 293T cells with pCMV_VSVG (Addgene plasmid no. 8454), psPax2 (Addgene plasmid no. 12260). Supernatant was collected day 2 and day 3 after transfection. Virus was concentrated with ultracentrifuge. Titer was measured with GFP percentage in limiting dilution infection to 293T cells. LARRY infection to human embryo models. LARRY lentivirus were infected to hEPSC in 6 -well plate during pre-differentiation stage in DMEM + 10% KoSR. MOI = 0.3 was used. Virus and cells were incubated for 2 hours at 37C, then washed twice with PBS and changed with fresh media. hEPSCs then underwent dissociation and aggregation to either hanging drop (HCEB) or Aggrewell (peri-gastruloids). Differentiation was conducted in corresponding media to HCEB or peri-gastruloids. Around 20-50 of human embryo models were harvested at each time point (Day 4, Day 6, and Day 12). The samples were dissociated and sorted with live cells with GFP+, and underwent 10X Chromium scRNA-seq.

### scRNA-seq sample preparation and sequencing

Preparation of single-cell suspensions. HCEB from D1 to D10 and human peri-gastruloids at D4, 6, 8 were harvested since EB formation in hanging drop.

The collected tissues were washed by 1× PBS, then quickly frozen and stored in liquid nitrogen. Nuclei extraction was separated by the mechanical extraction method ^51^. Firstly, put the tissues into 2ml Dounce homogenizer set and thaw them in homogenization buffer (containing 20mM Tris pH8.0, 500mM sucrose, 0.1% NP-40, 0.2U/μL RNase inhibitor, 1× protease inhibitor cocktail, 1% bovine serum albumin (BSA), and 0.1mM DTT. Use Dounce pestle A to grind the tissue 10 times, filter with 70μm cell filter, and then grind with Dounce pestle B 10 times, filter with 30um cell filter. Centrifuge at 500 × g for 5 minutes at 4°C to pellet the nuclei, and resuspend in the blocking buffer containing 1% BSA and 0.2U/μL RNase inhibitor in 1× PBS. Centrifuge again at 500 × g for 5 minutes and resuspend with Cell Resuspension Buffer (MGI).

Single-cell RNA library construction and sequencing. DNBelab C Series High-throughput Single-Cell RNA Library Preparation Set (MGI, #940-001818-00) was utilized for scRNA-seq library preparation. In brief, the single-cell suspensions were converted to barcoded scRNA-seq libraries through steps including droplet encapsulation, emulsion breakage, mRNA captured beads collection, reverse transcription, cDNA amplification, and purification. cDNA was sheared to short fragments with 300-500 bp, and indexed sequencing libraries were constructed according to the manufacturer’s protocol. Qualification was performed using the Qubit ssDNA Assay Kit (Thermo Fisher Scientific) and the Agilent Bioanalyzer 2100. All libraries were further sequenced by the MGISEQ-2000 sequencing platform with pair-end sequencing. The sequencing reads contained 30-bp read 1 (including the 10-bp cell barcode 1, 10-bp cell barcode 2 and 10-bp unique molecular identifiers (UMI)), 100-bp read 2 for gene sequences and 10-bp barcodes read for sample index.

### Single-cell lineage tracing data generation

Preparation of single-cell suspensions. Peri-gastruloids were harvested at D4, D6, and D8. HCEB organoids were harvested at D4 and D12 since EB formation in hanging drop. Organoids were washed with PBS, followed by mechanical chopping with scissors 20-30 times. The tissues were digested in 500 μL Accumax at 37 °C for 10-15 minutes and terminated with 500 μL PBS with 2% FBS. Cells were collected through the 40 μm cell filter. The cell suspension was centrifuged at 500 × g for 5 min and resuspended in FACS sorting buffer (1 × PBS with 2% FBS) for subsequent staining. Cell concentration was adjusted to around 3k cells per μL by counting with a hemocytometer.

Flow cytometry for scRNA-seq. Cells were stained in FACS sorting buffer with DAPI (BD Biosciences, catalog no. 564907) in 1:100 for 5 min at 4 °C. Cells were sorted by BD Influx flow cytometry equipment in CPOS at HKUMed. Cells were gated to exclude dead cells and doublets and collected in a chilled single-cell suspension medium (1x PBS with 0.04% BSA) for scRNA-seq library construction. Cell concentration was adjusted to around 500-1000 cells per μL counted with a hemocytometer.

scRNA-seq library preparation and sequencing. Single-cell encapsulation, library preparation, and sequencing were done at The University of Hong Kong, LKS Faculty of Medicine, Centre for PanorOmic Sciences (CPOS), Genomics Core. Single-cell encapsulation and cDNA libraries were prepared by Chromium Next GEM Single Cell 3’ Reagent Kit v3.1 and Chromium Next GEM Chip G Single Cell Kit. Around 8-16k live single cells of size 30 μm or smaller and of good viability were encapsulated, followed by reverse transcription and library preparation to harvest a pool of cDNA libraries. Libraries were sequenced using Illumina Novaseq 6000 for Pair-End 151bp sequencing. Individual samples had an average throughput of 100 Gb. We then retrieved the same library and amplified LARRY barcodes with PCR, followed by NGS.

### Construction of human and non-human primate embryonic developmental tree

Data collection and cleaning. Human published embryonic datasets were downloaded from E-MTAB-3929, E-MTAB-9388, GSE136447, GSE109555, GSA-Human: HRA005567^7,8,52–54^. Monkey embryo datasets were downloaded from public datasets ^55–59^. To balance the cell number at different embryo stages, the CS8 embryo sample was downsampled to 2,000 cells. Finally, altogether 7,823 cells from 149 human embryos were included. We adapted the original annotation from individual papers and hypoblast annotation ^60^ and merged some of the labels into higher-level groups.

Integration and batch correction. Samples from different human embryos were merged and processed as a single-cell experiment project. FastMNN was used to correct the batches, including the following steps: log-normalize the counts, get highly variable genes, compute the number of cells in each sample, reorder the samples by cell number, run fastMNN, and get the corrected PCs. After obtaining the corrected PCs, we used these corrected PCs to calculate the neighbors and generate the UMAP embedding.

### Single-cell RNA sequencing data processing

Single-cell RNA sequencing data processing from BGI platform (Alignment, Barcode Assignment, and UMI Counting). The sequencing data were processed using an open-source pipeline (https://github.com/MGI-tech-bioinformatics/DNBelab_C_Series_scRNA-analysis-software). Briefly, all samples underwent sample de-multiplexing, barcode processing, and single-cell 3’ unique molecular identifier (UMI) counting with default parameters. Processed reads were then aligned to the GRCh38 genome reference using STAR (2.5.1b). Valid cells were automatically identified based on the UMI number distribution of each cell by using the “barcodeRanks()” function of the DropletUtils tool to remove background beads and the beads that had UMI counts less than the threshold value. Finally, we used PISA to calculate the gene expression of cells and create a gene x cell matrix for each library.

Single-cell RNA sequencing data processing from 10X Genomics platform (Alignment, Barcode Assignment, and UMI Counting). The sequencing data were processed with CellRanger v8.0.1. Processed reads were then aligned to the GRCh38 genome reference. Gene x cell count matrix were obtained and used for further downstream analysis.

### Single-cell lineage tracing data analysis

Processing of LARRY barcodes. LARRY barcodes were extracted from an additional NGS sequencing library from a cDNA single-cell library. Sequencing reads mapping to the amplicon with the LARRY barcode were extracted from the raw sequencing reads using the fluorophore sequence GCTAGGAGAGACCATATGGGATCCGAT. The LARRY barcode was determined using the base pairs following the GFP sequence, given that the sequence matches the rules by which the LARRY barcode was constructed. Barcode extraction was performed using a modified version of the scripts provided in the original LARRY publication (https://github.com/AllonKleinLab/LARRY). Barcodes supported by fewer than five sequencing reads were discarded, and LARRY barcodes with a Hamming distance lower than three were merged. CoSpar algorithm was used to calculate the lineage probability with shared barcodes ^61^. X_Clone algorithm was used to pair lineage with shared barcodes ^62^.

### Data availability

Since NCBI lack of funding, the processed data and files generated in this study can be found on the download page of GitHub: https://github.com/CHAOYiming/hypoblast-blood.

### Code availability

Scripts for the data analysis have been deposited on GitHub: https://github.com/CHAOYiming/hypoblast-blood.

## Acknowledgement

We appreciate CPOS HKU for sequencing, flow cytometry, and imaging services. We particularly appreciate Hin Kwok’s expert input on library preparation and barcoding. We appreciate colleagues from the Centre for Translational Stem Cell Biology for constructive feedback and technical support. We appreciate funding from RGC GRF 17109424, RGC GRF 17115325, and InnoHK. We appreciate Patrick Tam, Bertie Göttgens, Tiago Rito, and Michael Hausser for their critical reading and advice on the manuscript, and Hugo de Jonge and Savani Anbalagan for their insights into hemoglobin-expressing cells.

## Author contribution

YC and RS designed the project and wrote the manuscript. YC conducted experiment design and informatics analysis. LL, HL, and ZT conducted embryo model experiments. SZ generated THP1 dCas9-VP64. XX, YX, DR and GL assisted embryo model experiments. SX generated *CDX2 -/-* EPSC. YH supervised informatics analysis. PL and RS supervised the project.

---

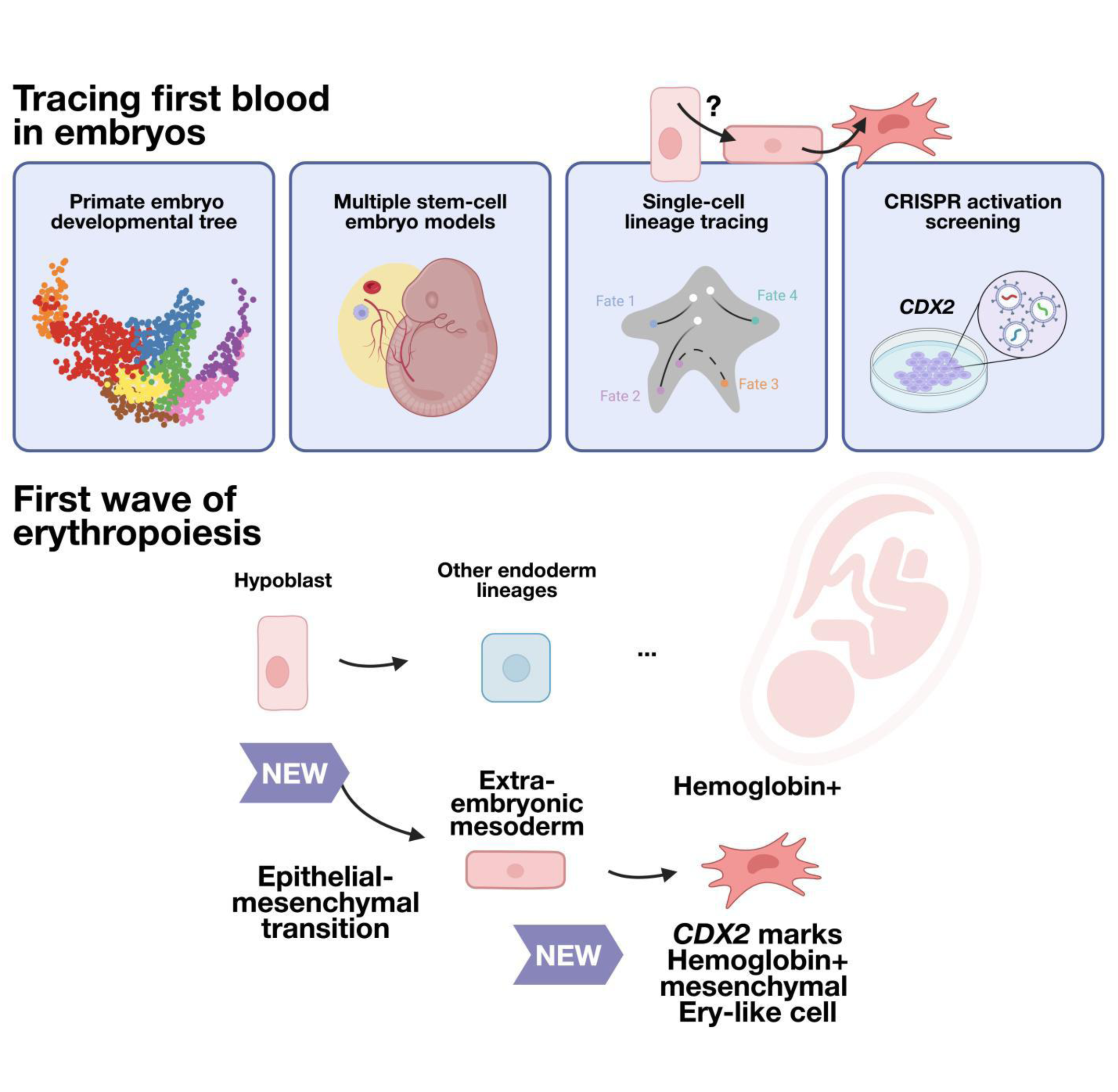

- Barcode-traced hypoblast fate in human embryo models;
- Hypoblast contributes to hemoglobin+ blood cells;
- CDX2 marks hypoblast blood that sustains oxygen supply in embryos before heart formation;
- Erythro-core regulatory network boosts erythropoiesis from human hypoblasts.

**Supplement Figure 1.**
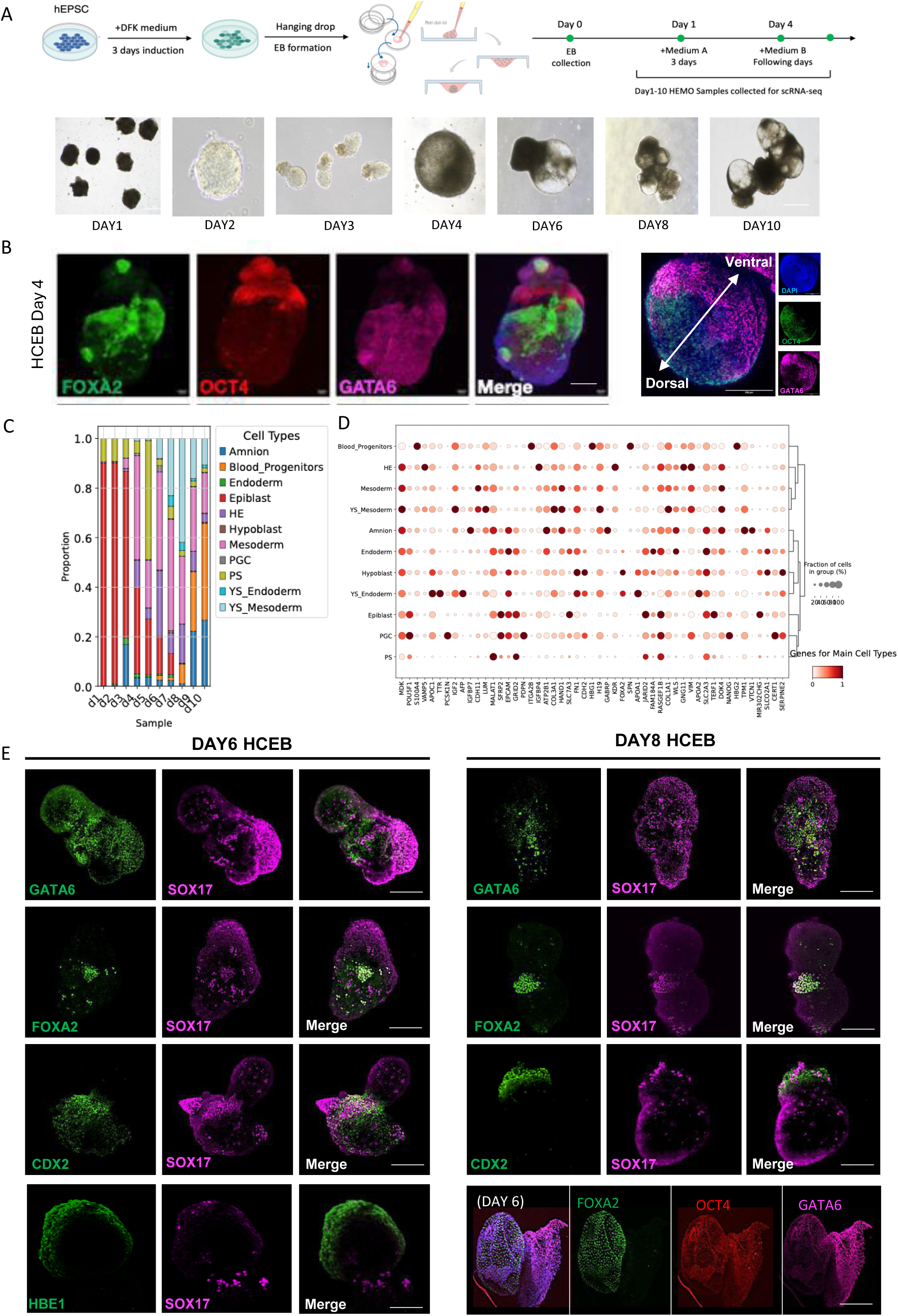
Human hypoblast comprises heterogeneous cell types and transitions to extraembryonic mesenchyme (corresponding to Figure 1). A. Hematopoiesis-competent embryoid (HCEB) generation from human expanded potential stem cells (hEPSC). HCEB was generated by preconditioning with KoSR media then step-by-step exposure to morphogens and cytokines. HCEB was cultured in 3D hanging drop. Scale bar = 200 um. B. Left: Epiblast (OCT4+) and hypoblast (FOXA2+, GATA6+) generation in Day 4 HCEB. Right: Epiblast (OCT4+)-hypoblast (GATA6+) polarization of Day 4 HCEB. Scale bar = 200 um. C. Cellular composition of HCEB at each time point scRNA-seq. D. Dot plot of marker genes used to define cell populations of HCEB. E. Immunofluorescence of HCEB at Day 6 and Day 8. Hypoblast markers GATA6, SOX17, and FOXA2 were partially overlapped by CDX2 and embryonic hemoglobin HBE1. Hypoblast heterogeneity indicated by reciprocal expression of FOXA2 (left) and GATA6 (right bottom). Scale bar = 100 um.

**Supplement Figure 2.**
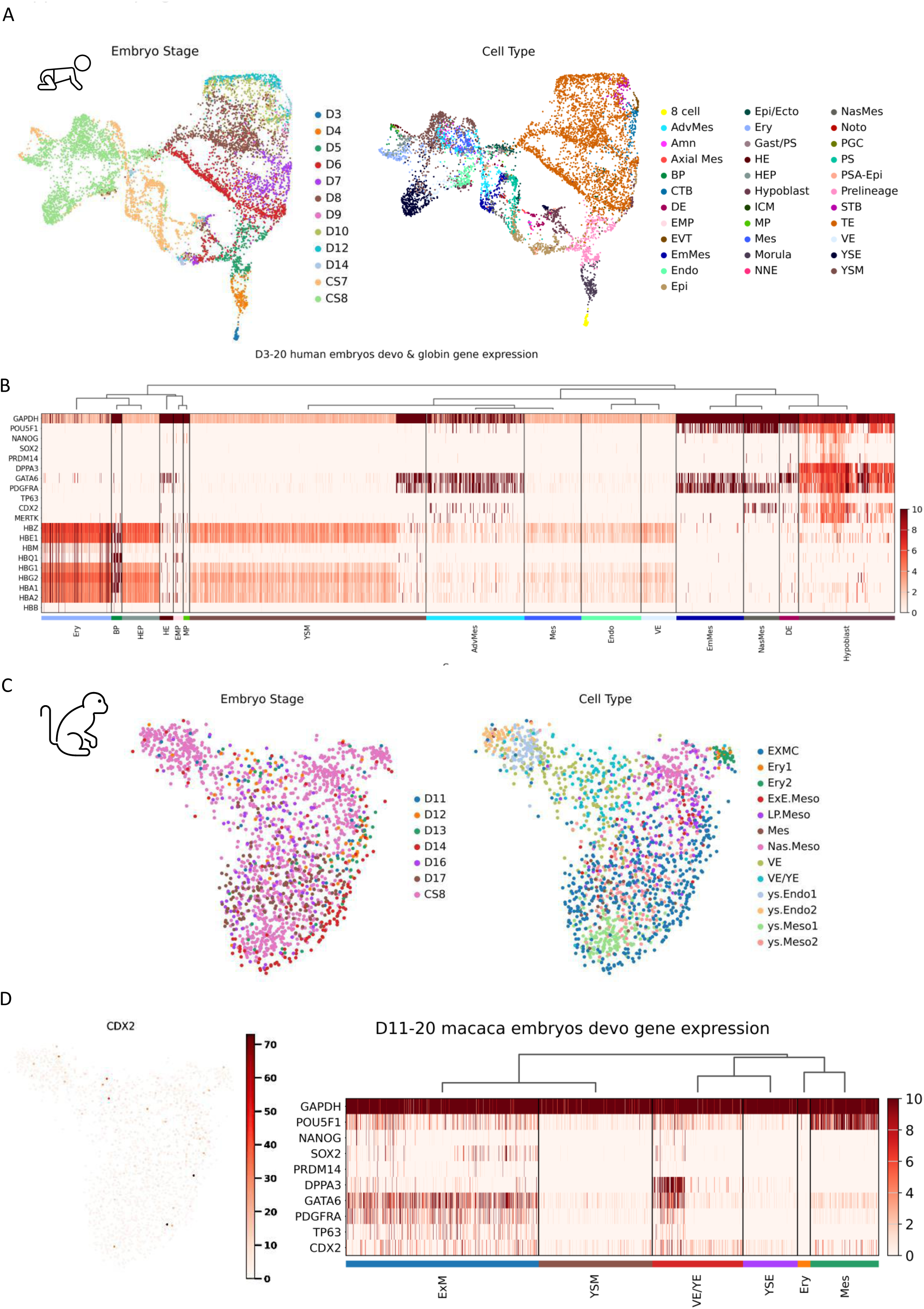
Hypoblast leads the lineage trajectory to extra-embryonic mesoderm and blood in human and non-human primate embryos (corresponding to Figure 2). A. A UMAP projection of the integration of five human embryonic datasets showing the construction of a human embryonic reference tree from blastocyst to the gastrula (D3 to CS8, equivalent to D20). The color of each data point represented the embryonic stage of each cell and cell identity. B. Heatmap showed the gene expression and the dendrogram of cell populations based on development-related markers in human embryos. CDX2 was expressed in hypoblast and ExM lineages. C. A UMAP projection of the integration of two Macaca embryonic datasets during gastrulation (D11 to CS8, equivalent to D20). The color of each data point represents the embryonic stage of each cell. D. CDX2 expression on UMAP of Macaca embryos. They were disperse, but expressed in ExM. The heatmap shows that CDX2 was expressed in hypoblast and ExM lineages in macaque embryos. A heatmap at the bottom showed hemoglobin expression in hypoblast and ExM lineages. AdvMes (advanced mesoderm); Amn (amnion); BP (blood progenitor); CTB (Cytotrophoblast); DE (definitive endoderm); EMP (erythro myeloid progenitor); EVT (extravillous trophoblast); EmMes (emerging mesoderm); Epi (epiblast); Ecto (ectoderm); HE (hemogenic endothelium); HEP (hematoendothelial progenitor); ICM (inner cell mass); MP (myeloid progenitor); NNE (nascent neuroectoderm); NasMes (nascent mesoderm); Noto (notochord); PGC (primordial germ cell); PS (primitive streak); STB (syncytiotrophoblast); VE (visceral endoderm); YSE (yolk sac endoderm); YSM (yolk sac mesoderm); EXMC (extraembryonic mesoderm); Ery (erythroblast); ExE (extraembryonic); ys (yolk sac); Endo (endoderm); Meso (mesoderm).

**Supplement Figure 3.**
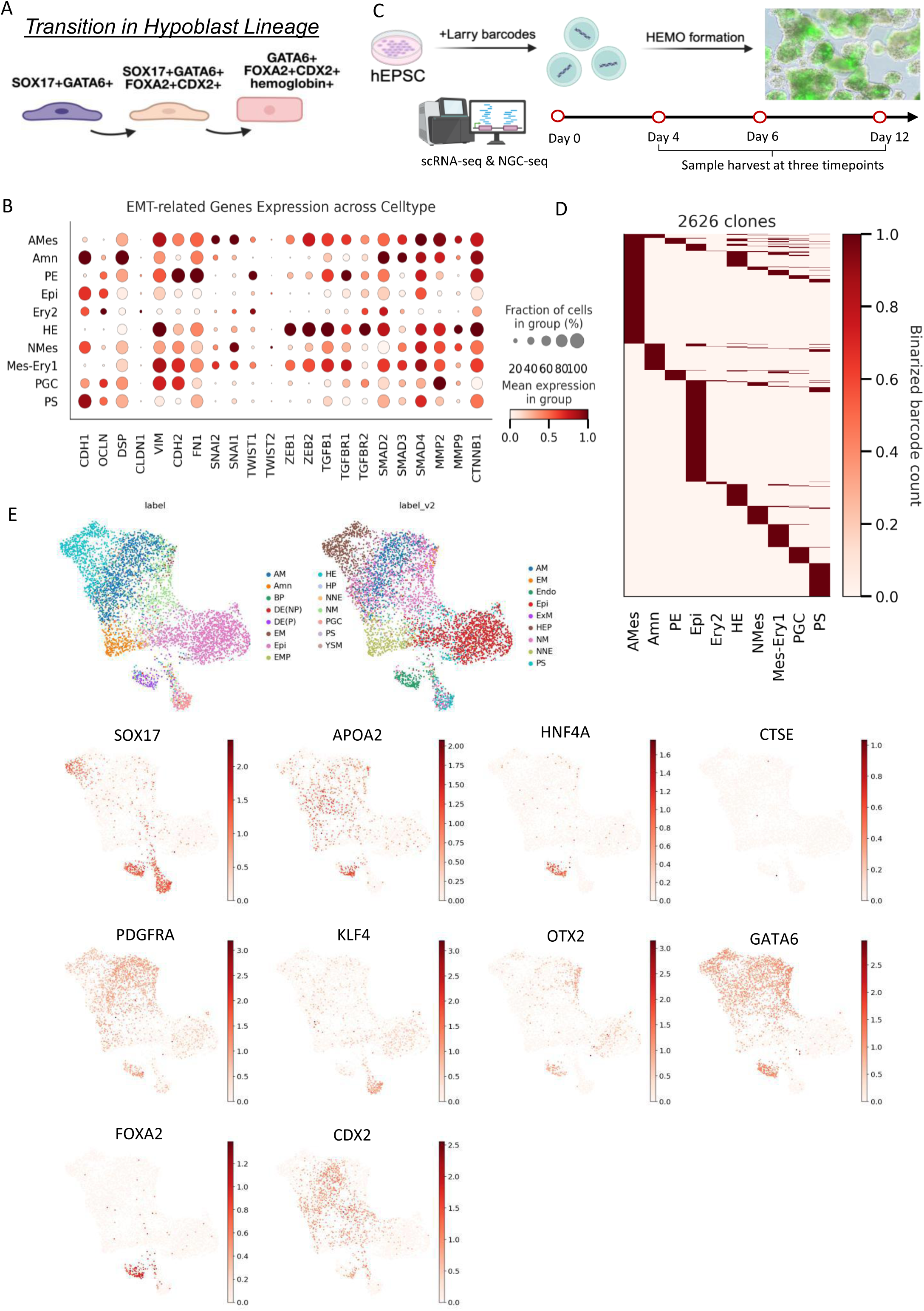
Hypoblast lineage trajectory (corresponding to Figure 3). A. Proposed model of human hypoblast lineage trajectory. SOX17+ GATA6+ cells (primary hypoblast) are followed by FOXA2+ CDX2+ cells (CDX2+ hypoblast), then hemoglobin+ cells (ExM). B. Key regulators of EMT, such as the SNAI family and ZEB family, were more highly expressed in the Mes-Ery population than in the Ery population. C. Experimental scheme of LARRY barcoding HCBE to trace hypoblast fate. LARRY was infected at hEPSC followed by HCEB formation. Samples were harvested at Day 4, 6 and 12, respectively. scRNA-seq and NGS-seq revealed cell populations and their corresponding barcodes. GFP indicated the LARRY infection in HCEB. D. NGS mapped each LARRY barcode to the cell population. 2626 clones were recovered. The heatmap showed the LARRY barcode distribution in each cell type. E. UMAP of Day 4 HCEB with hypoblast markers SOX17, APOA2, HNF4A, CTSE, PDGFRA, KLF4, OTX2, GATA6, FOXA2, and CDX2 were shown.

**Supplement Figure 4.**
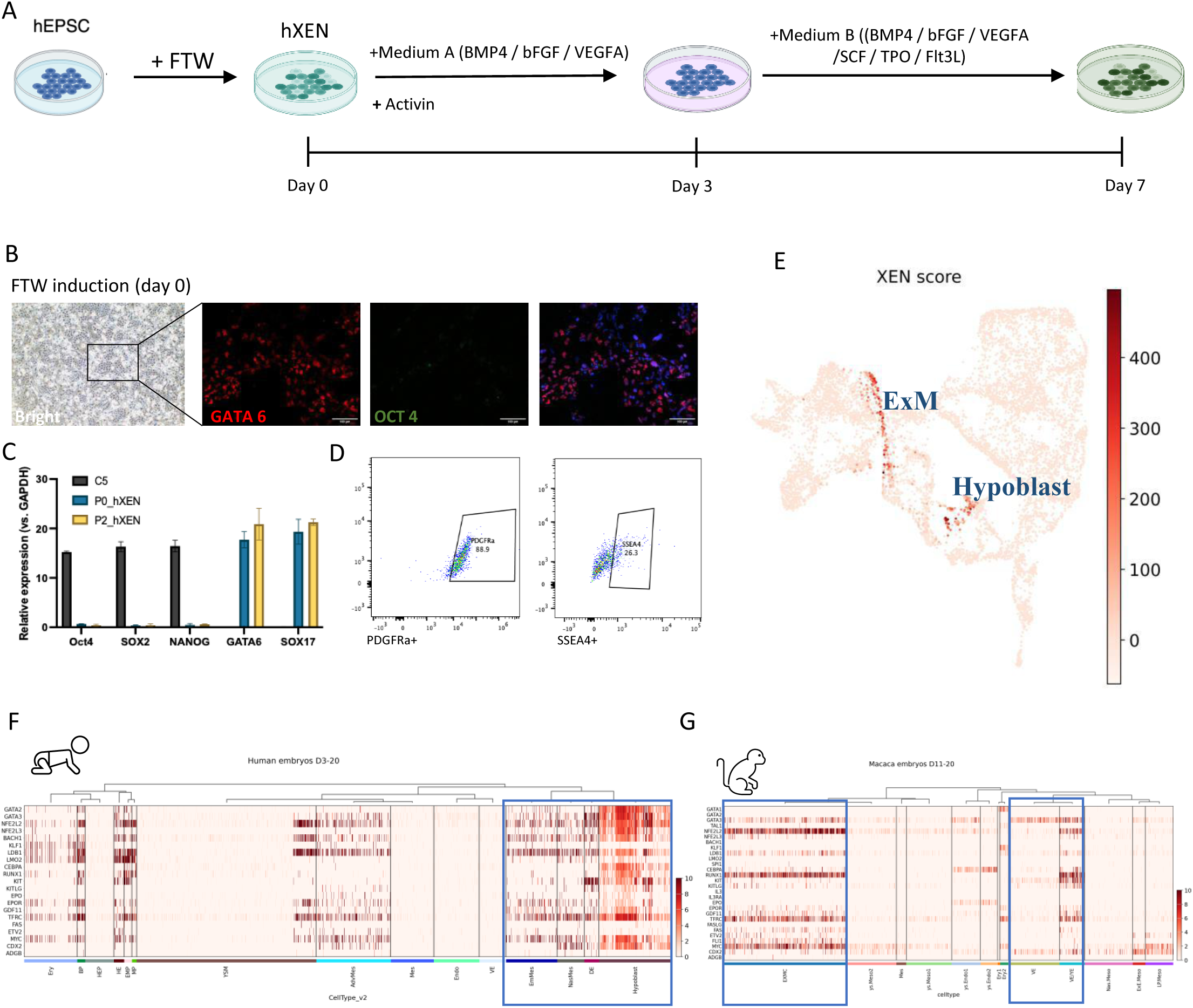
hXEN commits to hematopoietic lineage (corresponding to Figure 5). A. Establishment of human hypoblast cell line hXEN from human expanded potential stem cells with FTW media, followed by hemogenic culture over 7 days. B. Morphology of hXEN (left). hXEN expressed hypoblast marker GATA6 and was absent from pluripotent regulator OCT4 (right). Scale bar = 100 um. C. qRT-PCR of hXEN confirmed hypoblast markers GATA6 and SOX17 as well as a drop of pluripotent markers OCT4, SOX2, and NANOG. Data were shown with an error bar SEM. D. Flow cytometry of hXEN expressed hypoblast marker PDGFRa and reduced expression of pluripotent marker SSEA4. E. UMAP of human embryos showed the XEN (extra-embryonic endoderm) gene score of each cell. Hypoblast and extra-embryonic mesenchyme (ExM) had a significant XEN gene score. F. 30 erythropoiesis genes were expressed in hypoblasts and extraembryonic mesodermal cells in human embryo samples Day 3-20. G. 30 erythropoiesis genes were expressed in hypoblasts and extraembryonic mesodermal cells in macaque embryo samples Day 11-20.

